# Structure prediction of alternative protein conformations

**DOI:** 10.1101/2023.09.25.559256

**Authors:** Patrick Bryant

## Abstract

Proteins are dynamic molecules whose movements result in different conformations with different functions. Neural networks such as AlphaFold2 can predict the structure of single-chain proteins in the conformations most likely to exist in the PDB. However, almost all conformations in the PDB are seen during training. Therefore, it is not possible to assess whether alternative protein conformations can be predicted or if these are reproduced from memory. Here, we train a new structure prediction network on a conformational split of the PDB to generate alternative conformations. Our network, Cfold, enables efficient exploration of the conformational landscape of monomeric protein structures. 52% (81) of the nonredundant alternative protein conformations evaluated here are predicted with high accuracy (TM-score>0.8). Cfold is freely available at: https://github.com/patrickbryant1/Cfold

## Introduction

Structure prediction of single-chain proteins is highly accurate for ordered structures with AlphaFold2 [1] (AF), RoseTTAFold [2], ESMFold [3] and Omegafold [4]. Generating alternative protein conformations is another important problem which can inform function, but remains unsolved. Two complementary methods to generate alternative conformations with AF using multiple sequence alignment (MSA) clustering exist. The first uses a decreasing amount of sequence clusters [5] and the second a clustering procedure with DBscan [6] to generate diverse clusters [7]. These protocols are evaluated on very few structures (eight and six structures, respectively) that may be present in the training set of AF. Therefore, it is not known if AF generates these alternative conformations from memory.

Other methods to generate alternative conformations are based on molecular dynamics (MD) [8]. These try to sample the free energy landscape of a protein. However, there is no guarantee that this landscape contains any alternative conformations in its unaltered form. Most conformational changes in proteins occur due to physiological changes in the environment making one state more favourable than another [9,10]. Without knowing the necessary changes in the environment, it is not possible to model alternative conformations. In addition, MD requires vast computational resources and generally does not scale to larger proteins [11].

To improve upon the limited analyses previously performed and provide an answer to whether different conformations can be predicted, we extract a set of structures from the PDB [12] that have alternative conformations with substantial changes (a difference in TM-score [13] of at least 0.2 between structures) and are not homologous to the training set of AF. In total, we obtained 38 proteins with alternative conformations. We do not think this amount is sufficient to address the problem as the total number of structural clusters (TM-score≥0.8 within each cluster) is 6696, meaning that only 0.6% of possible structures would be evaluated.

Therefore, we create a conformational split of the PDB using structural clusters (TM-score≥0.8). We obtain 244 alternative conformations for evaluation which represent all sequences that have nonredundant structures that differ >0.2 in TM-score in the PDB. As AF (and other structure prediction methods) can’t be evaluated on this set due to having seen most of these conformations during training, we train a new version of AF on the conformational split which we name Cfold.

An important insight is that a network can, likely, not be trained to predict alternative conformations directly based on the coevolutionary information in an MSA. Firstly, there is too little data to properly develop and assess such a model. Although 244 alternative conformations may be enough for evaluation, it is too little for training (thousands are needed [1,14]). Secondly, a structure prediction network should learn to extract coevolutionary patterns that relate to a specific structure [15].

There may be many alternative conformations for a given protein and using the same coevolutionary representation created from an MSA should not result in different outputs. Therefore, the focus should be on generating different coevolutionary representations that represent different protein conformations, whatever these may be [16]. If an accurate mapping between a coevolutionary representation and a structure has been learned, applying the learned principles to different coevolutionary representations should result in different protein structures that represent a set of present alternative conformations.

## Results

### Predicting Alternative Protein Conformations

Many proteins occupy several different conformations, each one essential for the overall protein function. The information on different protein conformations is embedded in the evolutionary history of proteins and can be extracted through analysis of multiple sequence alignments (MSAs) [16]. However, one can not know beforehand what part of the full coevolutionary representation in an MSA relates to what protein conformation. Since a structure prediction network is tasked with predicting a single protein structure, obtaining descriptions of multiple conformations through an MSA poses a problem as the network has to pick one.

Here, we train a network to predict only one possible conformation using only sequence information (MSAs). We do this by dividing all single-chain structures in the PDB into structural clusters (Figure 1a). We then partition the identical sequences that are present in different clusters - the alternative conformations (Methods). Using one of the partitions of conformations, we train a structure prediction network similar to AlphaFold2 [1] and the remaining structural clusters of conformations are saved for evaluation.

**Figure 1.**
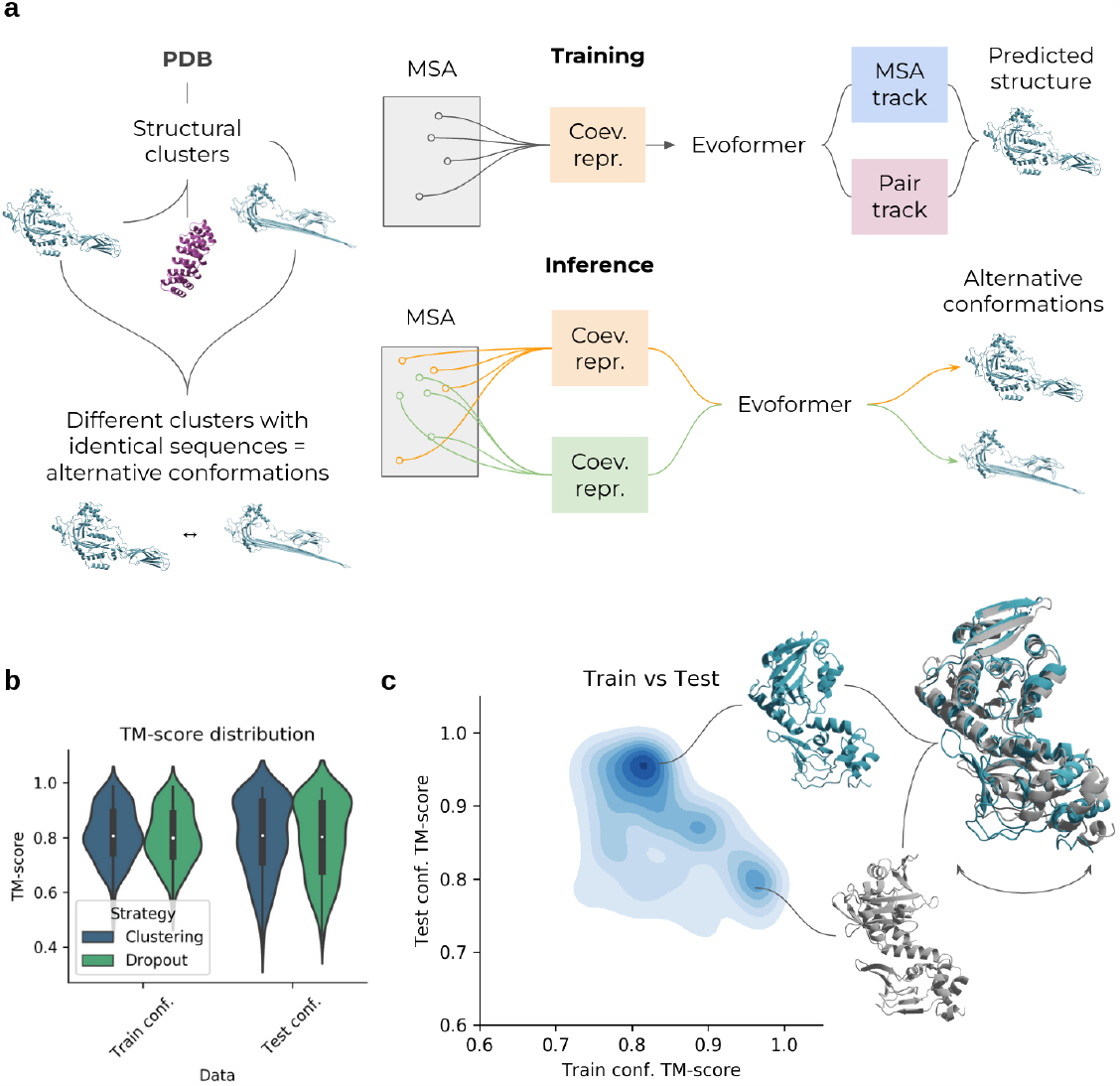
**a)** Description of Cfold. Using all monomeric protein structures in PDB we create a conformational split of structural clusters. Different conformations are defined as having >0.2 TM-score difference in identical sequence regions. A structure prediction network is trained on one partition of conformations and the remaining structural clusters of conformations are saved for evaluation. The network trained to predict structures is similar to the Evoformer of AlphaFold2 (Methods). Two tracks are present, one processing the multiple sequence alignment (MSA) representation and one the amino acid pair representation, the MSA- and pair tracks. At **training** time, one coevolutionary representation is created to predict one structure. At **inference** (the MSA clustering strategy is displayed here), the trained Evoformer network is used to predict alternative protein conformations by creating many different coevolutionary representations (orange/green). These are made by sampling and clustering different sequences from the full MSA. **b)** TM-score distributions of conformations trained and not trained on (test) for the different strategies (n=145 and 154 for dropout and MSA clustering, respectively). The MSA clustering results in slightly better results than using dropout. The best TM-score was selected for each method out of approximately 100 samples (Methods). **c)** Density plot of the TM-score to conformations in the training set vs. TM-score to unseen conformations using the best strategy (MSA clustering). The higher the density, the darker the colour. Only structures that could be predicted with a TM-score>0.8 for both train and test conformations among the samples taken are displayed (52 structures, n=5408 sampled predictions). Predicted structures corresponding to PDB IDs 4AVA/4AVB (blue/grey) are shown.

Training on structural partitions ensures that the network associates one input MSA with one protein conformation. Using the trained network, we then manipulate the statistical representation of the MSA to extract possible conformations described in the coevolution there. We evaluate the trained network on 155 nonredundant structures that can be predicted with the same folds when compared with conformations seen during training.

To find alternative conformations, we apply two different strategies (Methods):

1. Dropout [17–19]
2. MSA clustering [5]

Dropout is normally only used for training to make a network more robust. Using it at inference means that different information is excluded at random from each prediction, resulting in different outputs. The MSA clustering generates different coevolutionary representations by sampling different subsets of the MSA (Figure 1). This results in different inputs to the network, resulting in different outputs (alternative conformations).

MSA clustering proves to be slightly better compared to dropout, resulting in 81 of the alternative conformations being predicted with a TM-score >0.8 (52%, Figure 1b and Table 1) compared to 76 using dropout. Figure 1c shows a density plot comparing the TM-score of predicted structures towards the training set vs. unseen conformations (test set). The density is higher for unseen conformations at high TM-scores (>0.9), suggesting that the network can predict unseen protein conformations with high accuracy.

**Table 1.** Number of successful predictions (TM-score>0.8) for conformations in both train and test partitions. The total number represents the proteins that could be predicted out of the 155 evaluated (the remainder failed mainly due to RAM limitations, see Methods).

| Method | Train >0.8 | Test >0.8 | Train&Test partitions>0.8 | Total |
| --- | --- | --- | --- | --- |
| MSA clustering | 81 | 81 | 52 | 154 |
| Dropout | 72 | 76 | 45 | 145 |

### Types of conformational changes

Not all conformational changes are equal. Some are more substantial than others, resulting in e.g. fold switches (alpha helices turned to beta sheets or vice versa) while others result from a movement of a single domain around a “hinge”. We categorise conformational changes into three types (Figure 2):

**Figure 2.**
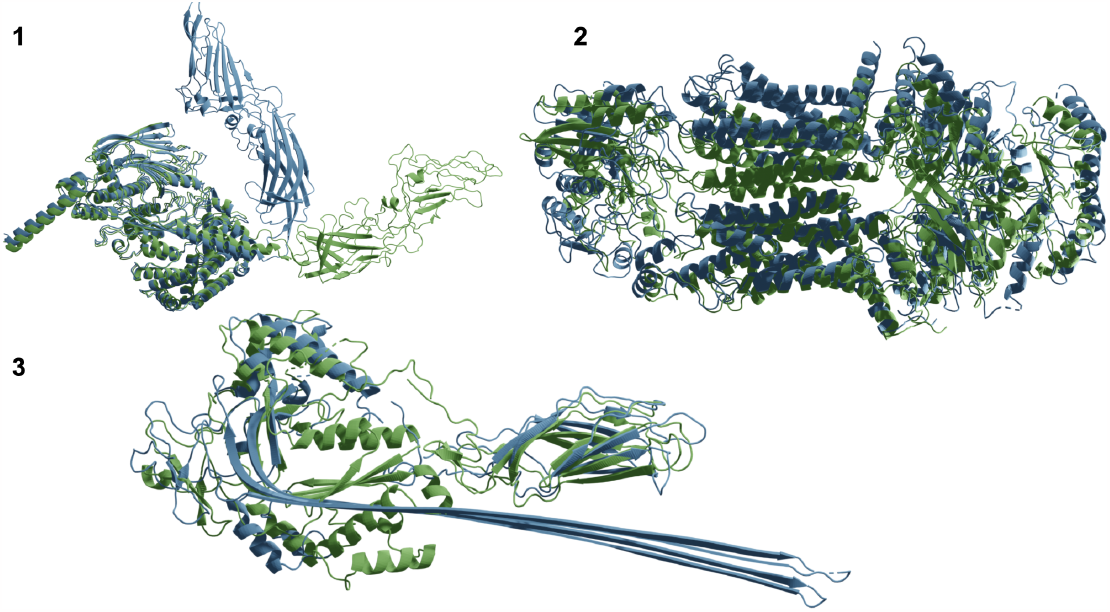
Different types of conformational changes; hinge motions (1, PDB IDs 7WHM/7WHN), rearrangements (2, PDB IDs 7W01/7W02) and fold switches (3, PDB IDs 3NSJ/7PAG). The most frequent conformational change is the rearrangement (n=180) followed by the hinge motions (m=62) and the fold switches are rare (n=3).

1. Hinge motion: The structure of all domains is intact and the change is the relative orientation between the domains.
2. Rearrangements: The domains change, but the secondary structure elements remain the same.
3. Fold switches. This is a rare type of conformational change where alpha helices turn to beta sheets or loops or vice-versa.

In total, there are 63 hinge motions, 180 rearrangements and 3 fold switches in the PDB fulfilling the criteria set for alternative conformations here (TM-score difference>0.2 across identical sequence regions). Among the structures that can be predicted with a TM-score≥0.6 (155 structures, Methods) there are 40 hinge motions, 114 rearrangements and 1 fold switch.

### Structural change and accuracy

The structural change (TM-score btw conformations) vs. the prediction accuracy (TM-score) for the test conformations can be seen in Figure 3a. The accuracy decreases with the structural change, even though there are accurate predictions at changes as large as 0.4 TM-score. Hinge motions are less accurately predicted than rearrangements and fold switches (Figure 3b). All three types of conformational change can result in both larger and smaller overall structural changes (Figure 6c), suggesting that the type of change does not determine the accuracy but the amount of structural change does.

**Figure 3.**
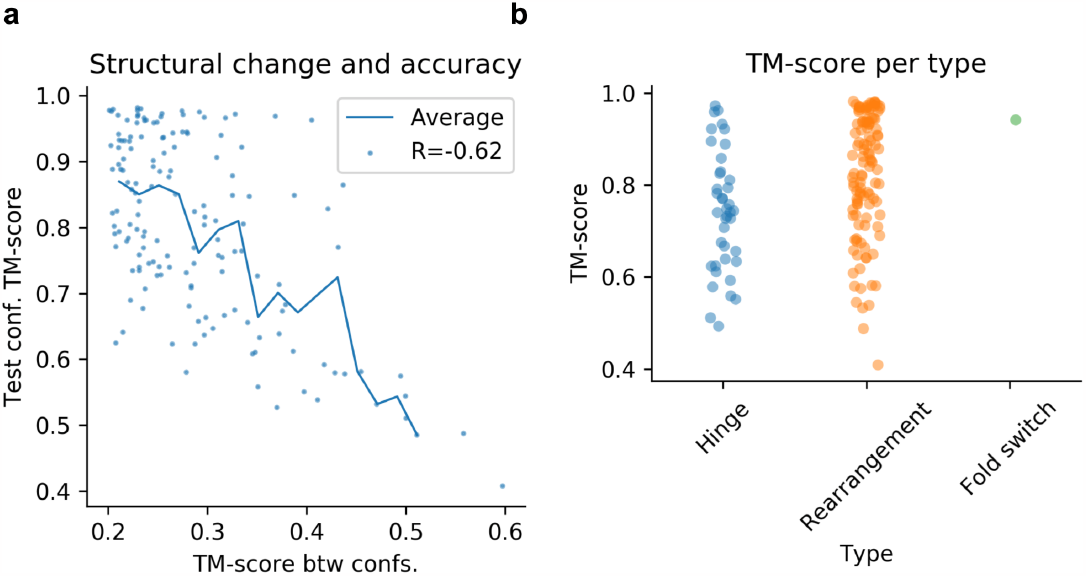
**a)** Structural change and accuracy. The x-axis shows the TM-score difference between the native conformations and the y-axis is the TM-score for the prediction and the structure of the test confirmation. The Pearson R is -0.62 suggesting that there is a relationship between the structural change and accuracy and that structures that change more between their conformations are harder to predict. The points represent each structure (n=155) and the line is the running average using a step size of 0.02 TM-score. **b)** Conformational type and accuracy. The hinge motions are less accurately predicted than the rearrangements and fold switches. This is likely explained by the findings in a.

### Selecting accurate conformations

The predicted lDDT (plDDT [1]) does not reflect the TM-score towards different conformations (Pearson R=0.52, Figure 4a). At the same time, the plDDT score is highly correlated to the overall structural accuracy on the validation set when not considering alternative conformations (Pearson R>0.9, Figure 9c). In the case of alternative conformations, it seems that the plDDT is rather a metric for how the structure may move, describing protein flexibility.

**Figure 4.**
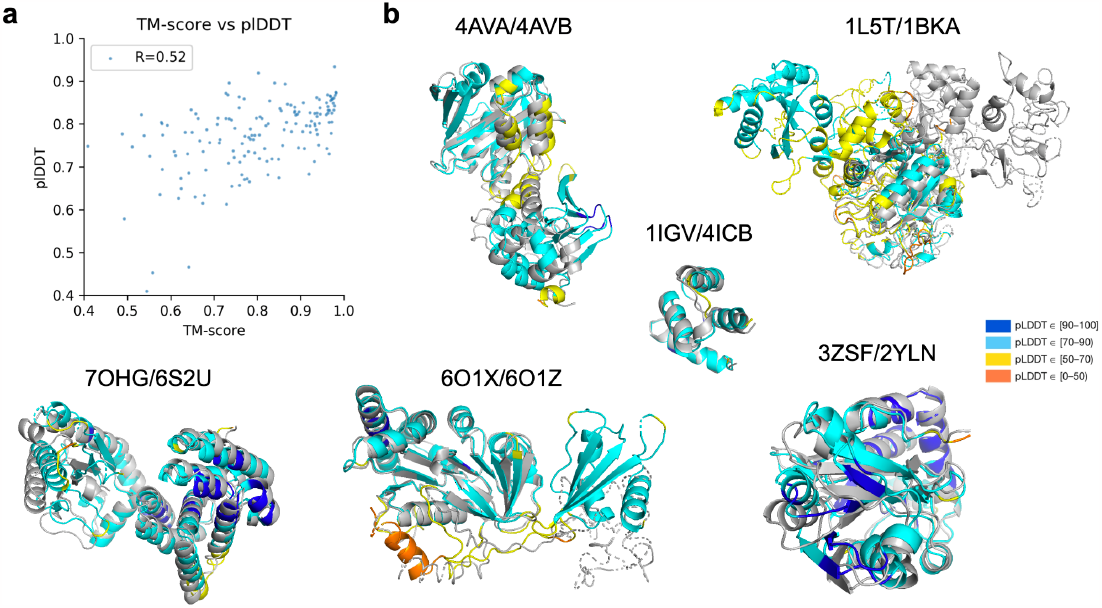
**a)** TM-score vs plDDT. The Pearson correlation between the TM-score and plDDT is low (0.52), suggesting that the quality metric can’t be used to distinguish between alternative conformations. **b)** Examples of predicted conformations coloured by plDDT/grey and in structural superposition. Some predictions are less accurate than others, resulting in chain breaks and low plDDT regions (orange).

The regions with lower plDDT tend to be flexible regions where the conformations change (Figure 4b). This seems to be the case for predictions for PDB IDs 4AVA/4AVB, 1L5T/2BKA and 7OHG/6S2U. For 6O1X/6O1Z, one of the sampled conformations has low plDDT in the region where the conformations differ and the other high. For 3ZSF/2YLN, both conformations have high plDDT overall and no flexible central region as in 4AVA/4AVB is found. This suggests that the plDDT is to be related to the regions that may change between conformations in some cases, but not in all.

### Rescuing failed predictions with increased sampling

To see if it is possible to “rescue” failed predictions by increasing the number of recycles and samples, we select 10 examples that are predicted with TM-score<0.8 at random. We increase the number of recycles to 10/20 and take 50/100 samples per cluster size with the MSA cluster procedure (Methods). Figure 5a shows the previous best TM-scores towards the test conformations vs. the best with the increased sampling and recycling. Increasing the number of recycles or samples has a negligible effect on the outcome, with only one of the targets displaying an improvement to a TM-score>0.8 and all scores are only moderately improved.

**Figure 5.**
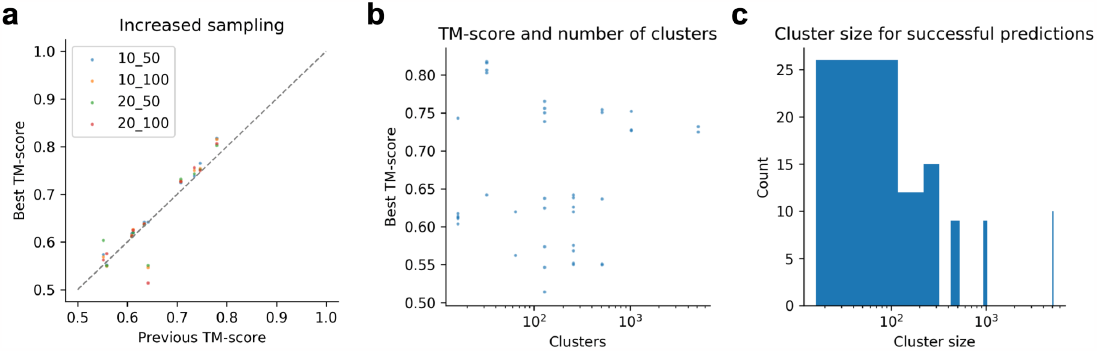
**a)** The best TM-scores obtained previously using 3 recycles and 13 samples per cluster size vs. 10/20 recycles and 60/100 samples per cluster size. Ten structures from the test set that were unsuccessful (TM-score<0.8) were used here with PDB IDS: 2NRV, 4NTJ, 5WU4, 4WXX, 6WBO, 5LJ8, 1M61, 4WTV, 2E1R and 6YHK. Increasing the recycles or number of samples has a negligible effect. **b)** TM-score and number of clusters for the rescue set. Using more clusters does not seem to be beneficial. **c)** MSA cluster size distribution for the best TM-scores towards the test set using the successful predictions (n=81, Table 1). Using more clusters does not appear to be beneficial in most cases as most high scores are found at cluster sizes <100.

Figure 5b shows the best TM-score and the number of clusters used to obtain this in the rescue attempt. There is no relationship with cluster size and the only successful example is obtained at a small cluster size of only 32 sequences. Therefore, there is no point in exploring the more expensive sampling settings with thousands of sequences, more recycles or samples. Figure 5c shows the distribution of cluster sizes using the best TM-scores towards the test set for the successful predictions (n=81, Table 1). Increasing the number of clusters does not appear to be beneficial in most cases as most high scores are found at cluster sizes <100.

## Discussion

### Structure prediction of alternative conformations with Cfold

The purpose of Cfold is not to replace AlphaFold2 in the prediction of protein structures, but to instead predict more than one conformation of a structure when this is possible. These *conformational ensembles* represent multiple important states of proteins that can inform functional aspects that a single structure can not. Currently, protein structure is viewed as a single snapshot in time while in reality, it should be viewed more like a video. This video is the continuous collection of states that make up the protein and its functions.

Cfold predicts 81/155 unseen structures with high accuracy (TM-score>0.8). Here, 13 samples and 3 recycles were used per cluster size, resulting in approximately 100 samples (Methods). However, it has been found that over 20 recycles and thousands of samples are necessary to obtain accurate predictions in challenging cases for multimeric proteins [17]. To see if this may be the case for the single-chain proteins evaluated here as well, we increased the recycles to 10/20 and took 50/100 samples per cluster size. We see no significant improvement in the scores. Using a higher number of clusters for the predictions is not beneficial either, suggesting that it is best to run the cheaper settings with fewer recycles, fewer samples and a lower number of clusters.

Proteins with larger structural changes are harder to predict and at changes >0.4 TM-score, the accuracy is poor. This is a limitation of the network and new methods that do not rely on coevolutionary information may be needed to capture these drastic changes. The type of conformational change does not seem to impact the results, suggesting that if the structural difference between conformations is in the range of 0.2-0.4 TM-score, there is a good chance of obtaining accurate predictions of alternative conformations (57%, 78/138).

### Coevolution and conformational changes

Conformational changes are direct effects of environmental changes. Therefore, it is not possible to capture them using e.g. molecular dynamics which samples structures in their equilibrium states, assuming that this process is accurate. Specific knowledge of the exact environmental changes is necessary and we see no realistic possibility of constructing a procedure to enumerate these as anything (including unknown phenomena) may happen in the cell. The reason that conformational states can still be predicted is that they will leave distinctive traces throughout evolution. By extracting these coevolutionary patterns from MSAs it is possible to also extract different conformational states without knowing what induces them.

### Applications and extensions

As Cfold can predict alternative conformational states with high accuracy in the majority of cases, it is possible to study alternative conformations using available protein sequence data and validate these with experiments for structure determination. We hope that Cfold will aid in extending known alternative conformations as the current pool of 244 conformations out of the 10116 sequence clusters (2.4%) suggests that there may be many unknown states of proteins. Alternatively, it must be rare for monomeric structures to take on alternative conformational states which poses questions of if dynamical studies are then meaningful for most proteins.

We expect that Cfold will be useful for a variety of researchers in elucidating various protein mechanisms resulting from conformational changes. To this end, we have made Cfold available through an online application (Google Colab) so that researchers with limited computational knowledge can also apply our method. We aim to extend Cfold to multimeric structures, where more alternative conformations are expected based on varying interaction partners and stoichiometries.

## Methods

### Proteins with alternative conformations not present in the AlphaFold2 training set

AlphaFold2 (AF) was trained on all structures in the PDB[12] with a maximum release date of 30 April 2018. To analyse if AF can predict alternative conformations it is necessary to first obtain structures with conformations not present in this training set. We obtained such a set by:

1. Selecting all monomeric protein structures from the PDB on 2023-07-05 determined by X-RAY diffraction or Electron Microscopy (EM) with a resolution ≤5 Å. We excluded NMR structures as the flexibility in these may give rise to the appearance of alternative conformations when there are none. In total, 69861 structures were obtained. 51183 were deposited on or before 30 April 2018 and 18678 after.

2. We extracted the first protein chain in each PDB file and the corresponding sequences with less than 80% non-regular amino acids and more than 50 residues. 68953/69861 proteins fulfilled these criteria (99%).

3. We clustered the sequences at 30% identity using MMseqs2 (version f5f780acd64482cd59b46eae0a107f763cd17b4d) [20]. This is a more stringent cutoff than the 40% identity used in AlphaFold2[1] to ensure that similar structures are indeed captured within the clusters.

`mmseqs easy-cluster examples/DB.fasta clusterRes tmp --min-seq-id 0.3 -c 0.8 --cov-mode 1`

4. This resulted in 5452 clusters with more than one entry (10116 clusters in total). From the sequence clusters with over one entry, we used TM-align (version f0824499d8ab4fa84b2e75d253de80ab2c894c56) [13] to perform pairwise structural alignment of all entries within each cluster. We created structural clusters if the pairwise TM-score≥0.8. 64208 out of the 68953 structures (93%) could be clustered into 6696 structural clusters.

5. We now checked for sequence clusters that have different structural clusters (different conformations) before and after April 30 2018. We obtained 900 putative alternative conformations.

6. To see if these are truly alternative conformations and not a result of sequence variations, we clustered the sequences again on 90% identity using MMseqs2:

`mmseqs easy-cluster examples/DB.fasta clusterRes tmp --min-seq-id 0.9 -c 0.8 --cov-mode 1`

7. We obtained 14662 clusters out of 64208 sequences in total. 7806 of the clusters have more than one member and 64 maps to alternative conformations (some with as many as 9 different ones).

8. To see if these are truly alternative conformations and not a result of length differences or low resolution, we performed a manual check. We find that some putative alternative conformations are a result of variations in N/C-terminal loops, mutations or disconnected chains (breaks) and such structures were excluded. The manual check resulted in 38 alternative conformations in total.

### Proteins with alternative conformations in the PDB

To assess all proteins with alternative conformations from the PDB, we continue from step 4 above using the structural clusters at 0.8 TM-score. We skip step 5 and go directly to step 6 (as no date cutoff is necessary here) to see if these are truly alternative conformations and not a result of sequence variations. **Figure 6** shows the entire data selection workflow starting from step 1 above.

- After step 6 (clustering at 90% sequence identity), we analyse all sequence clusters at 90% (14662 clusters) that contain at least two different structural clusters. In total, there are 627 such sequence clusters.
- To reduce the number of necessary manual checks, we do pairwise sequence alignment of all sequences within each structural cluster of the 90% sequence clusters. Using these sequence alignments, we extract the corresponding structural areas and perform another structural comparison with TM-align. Only if the structures are still different (TM-score<0.8), do we consider them to be putative alternative conformations.
- We obtain 311 such examples that are within the same 90% sequence identity cluster but have aligned sequence regions with corresponding structures that differ >0.2 in TM-score. We select representatives with the longest sequence overlaps and the biggest TM-score difference from each of the structural clusters. We perform a manual check to ensure that the conformations are indeed different and not a result of variable loop regions, low resolution or other experimental differences.
- We obtained 245 sequence clusters at 90% with alternative conformations (TM-score difference>0.2) belonging to 458 different structural clusters. Only one of the conformations of these is selected for training and the remainder is saved for evaluation. We select the clusters at random. In total, there are 6474 structural clusters composed of 59946 sequences for training and 222 composed of 4263 sequences for evaluation with a total of 244 alternative conformations (some structural clusters are present in more than one alternative conformation). We select 315 of the training structural clusters at random for validating the structure prediction performance (5%, 3539 sequences).

**Figure 6.**
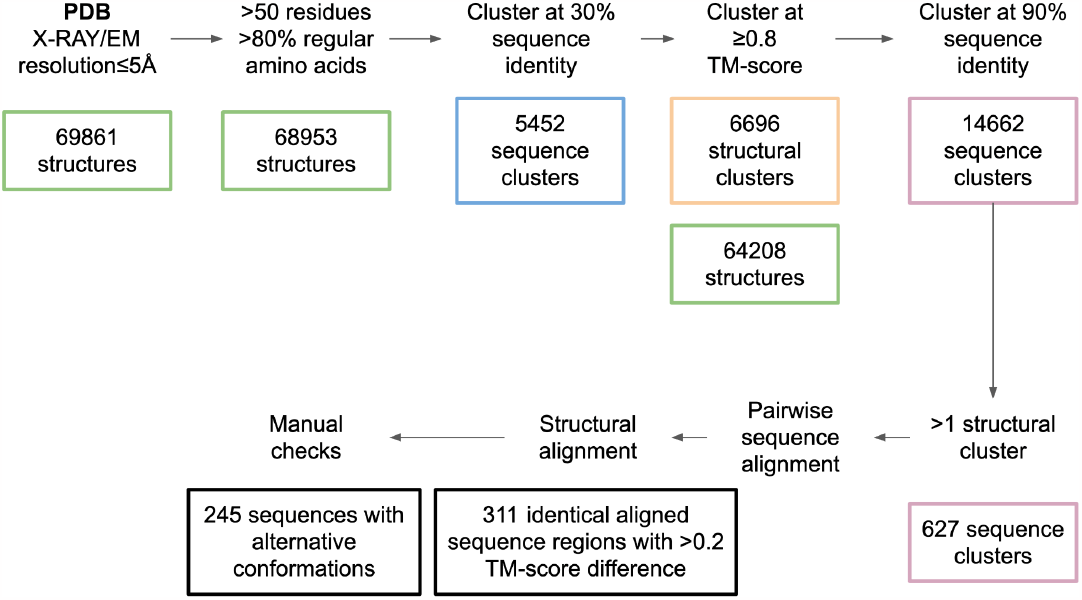
All single-chain structures with at least 50 residues and 80% regular amino acids determined by cryo-EM or X-ray crystallography with a resolution ≤5Å were selected. These were then clustered at 30% sequence identity and 0.8 TM-score. Within the structural clusters, sequences with 90% sequence identity in different structural clusters were analysed manually to annotate alternative conformations. In total, 245 such sequences were found.

Most structures with alternative conformations are below 500 residues and the structural change between conformations is 0.2-0.3 TM-score (Figure 7). The most represented type of conformational change is rearrangements, and the least fold switches (Figure 7c). In total, there are 62 hinge motions, 180 rearrangements and 3 fold switches.

**Figure 7.**
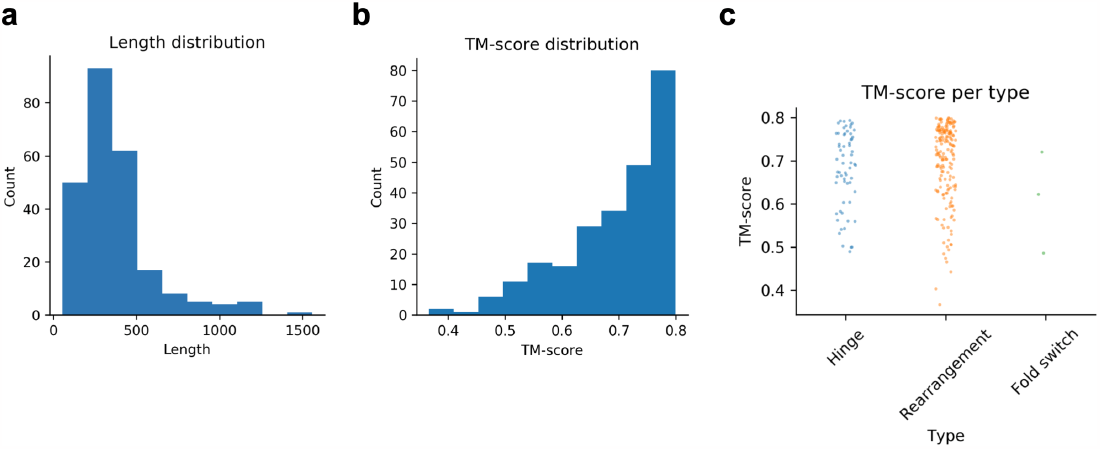
**a)** Sequence length distribution. Most sequences are below 500 residues. **b)** TM-score distribution between alternative conformations **c)** Strip plot of TM-score between alternative conformations per type (n=62, n=180 and n=3 for hinge motion, rearrangement and fold switch, respectively). The fold switches are fewer with lower TM-scores than the hinge motions and rearrangements.

### Training, validation and test partitions

Table 2 displays the number of sequences and structural clusters in each partition. Approximately 5% of the sequences and structural clusters were used for validation and testing and 90% for training.

**Table 2.**
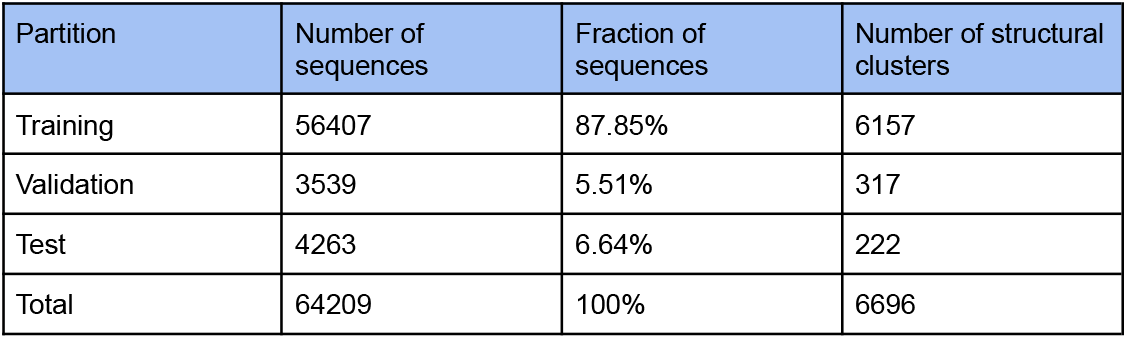
Number of sequences and structural clusters in the training, validation and test partitions. The procedure for selection and generation of these is outlined in “Proteins with alternative conformations in the PDB”.

### Scoring

TM-score from TM-align [13] was used for all scoring. We consider conformational changes to have at least a difference of 0.2 in TM-score. This is because smaller changes are very difficult to distinguish. Note that having a TM-score above 0.5 is generally considered as having similar folds [21] and this analysis is therefore very refined (a TM-score above 0.8 is highly accurate).

Another reason is the evaluation of predicted structures. If a change of only 0.1 TM-score is permitted, the predictions have to have a score of >0.9 to be able to distinguish between the different conformations. This is very unlikely as this means that almost all atoms are coincidental between predicted and native structures. Therefore, we use a threshold of 0.2 TM-score.

### MSA generation

AlphaFold2 generates three different MSAs. This process constitutes the main bottleneck for the predictions as very large databases such as the Big Fantastic Database [22,23] are searched which is very time-consuming [24]. To simplify this process we instead search only uniclust30_2018_08 [25] with HHblits (from HH-suite [26] version 3.1.0):

`hhblits -E 0.001 -all -oa3m -n 2`

We note that it is not the number of hits or the number of effective sequences [27,28] in the MSAs that determine the outcome, but rather the coevolution present [16]. This is an elusive concept extracted by the network and it is possible to improve upon the MSA generation at inference by searching larger databases. Indeed, AF itself has been improved upon by searching larger sets of metagenomic sequences[29].

### Structure prediction network

#### Architecture

The structure prediction network trained here is almost a complete copy of AlphaFold2[1] for monomeric structure prediction. The main difference in the architecture is that the template track - which processes similar structures - is removed here to focus on the MSA only.

Templates can capture alternative conformational states, although this would mean that these states are not predicted but instead copied from these templates.

The configuration for all layers and modules is the same as in AlphaFold2 (Table 3).

**Table 3.** Number of blocks and cluster sizes used in the network.

| Component | Size |
| --- | --- |
| Evoformer blocks | 48 |
| MSA clusters | 128 |
| Extra MSA clusters | 1024 |
| Structure module blocks | 8 |

#### Training

The main difference overall between the network here (Cfold) and other structure prediction networks is how it is trained. We train Cfold on structural clusters of sequences which enables learning a one-to-one mapping of a given MSA and a protein conformation. The structural clusters used for training are selected at random, which means that the network has to learn to extract specific coevolutionary information from each MSA that relates to a certain conformation that is learned during training.

This sets the network up to be more focused on one structural description in the MSA, important for the possibility of later evaluating the impact of different MSAs on the outcome. We sample the structural clusters with inverse probability to the cluster size. This allows the network to learn a more refined mapping from each sequence to the exact structure compared to using sequence clusters alone.

The effective batch size is 24 distributed across 8 NVIDIA A100 GPUs (3 examples per GPU) with a crop size of 256 residues. We apply the same losses as in AlphaFold2:

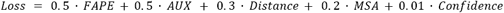

Where *FAPE* is the frame aligned point error, *AUX* a combination of the *FAPE* and angular losses, *Distance* a pairwise distance loss, *MSA* a loss over predicting masked out MSA positions and *Confidence* the difference between true and predicted lDDT scores. These losses are defined exactly as in AlphaFold2 and we refer to the description there [1].

We use a learning rate of 1e^-3^ with 1000 steps of linear warmup and clip the gradients with a global norm of 0.1 as in AlphaFold2. The optimiser is Adam [30] applied through the Optax package in JAX which the entire network is written in (JAX version 0.3.24, https://github.com/deepmind/jax/tree/main). The model is trained until convergence with a total of 74000 steps (compared to 78125 in AlphaFold2). The different losses and how they converge can be seen in Figure 8. During training, 56345/56407 (99.9%) of the training examples could be loaded due to file system issues blocking access to some data. We do not expect that this will impact the performance of the network.

**Figure 8.**
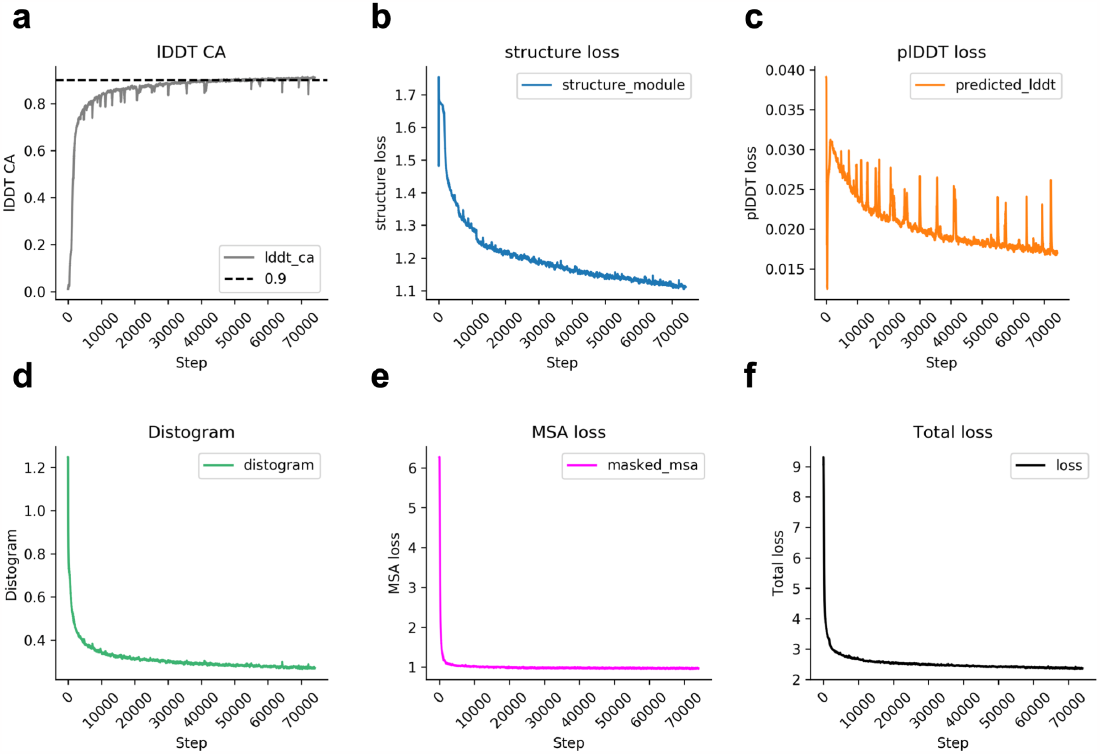
Train losses and metrics vs. training steps. The losses have been smoothed with an exponential moving average (step size=100). The lDDT CA increases continuously, although almost all performance is reached in the first 10000 steps (a). First, the MSA (e) and distogram losses (d) saturate, followed by the structural module loss (b) and plDDT loss (c). The total loss (f) is the sum of distogram, MSA, predicted lDDT (plDDT) and structural module losses.

### Structure prediction validation

To validate the training of Cfold, we use the full-length structures taken from 315 structural clusters (one per cluster sampled randomly) and 3 recycling iterations. From the 315 sampled structures, 307 can be evaluated (97%) due to structural inconsistencies (e.g. missing CAs) causing errors with TM-align and the lDDT calculations.

**Figure 9a** shows the TM-score distribution vs. the training step and **b** the lDDT scores. The best model weights are obtained at **10000 steps** and these are the ones used in Cfold. It takes approximately 3 days to train the model to this point. The relationship between the predicted and true lDDT scores remains stable throughout the training process (**Figure 9c**). The median TM-scores at 20000 and 30000 steps are similar to those of 10000, but at ≥10000 steps the secondary structure of the beta sheets is inaccurate (**Figure 9d**).

**Figure 9.**
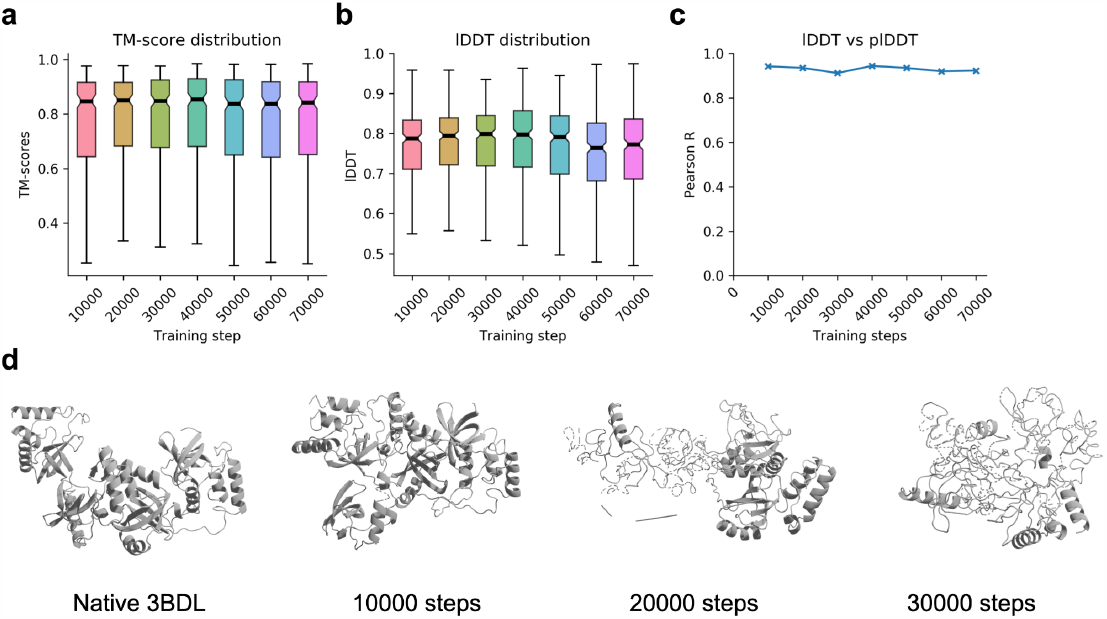
**a)** TM-score distribution for the validation set vs. training step (n=307). **b)** lDDT CA distributions vs. training step (n=307). **c)** Pearson correlations for lDDT vs. plDDT for the alpha carbons (CA, n=307). The correlation coefficients (R) remain close to 0.9 throughout the training. **d)** Predicted structures for PDB ID 3BDL from the validation set at different training steps. The TM-scores are 0.59, 0.53 and 0.55 for 10000, 20000 and 30000 steps, respectively. Visually, one can see that the network starts to make worse predictions >10000 training steps, although this is not apparent from the metrics in a,b and c. This suggests that the network starts to overfit to certain structures at an early stage.

#### Structural limitations

To correct for bond length violations and overlapping atoms, specific losses are added during the fine-tuning stage in AlphaFold2. In addition, one can obtain slightly more accurate structures by relaxing predictions in the Amber force field [31]. Here, we did not perform fine-tuning as the main objective is to sample globally different conformations. To generate highly accurate structures locally, the same Amber force field can be applied to obtain consistent side chain angles for all amino acids. We did not relax the structures here as this will likely not impact the overall performance.

### Conformational generation

We select structures whose conformations are in the train set from the test clusters that Cfold can predict with a TM-score of at least 0.6 using one prediction (0.5 is considered to be of a similar fold [cite]), resulting in 155 out of 234 possible structures (66%). We take the minimum TM-scores from the TM-align structural superpositions here. To analyse if Cfold can predict the structure of alternative conformations of these, we apply two different strategies:

1. Dropout [17–19]
2. MSA clustering [5]

For the dropout, we simply activate dropout everywhere except for in the structural module following AFsample[17] and make 100 predictions in total. For the MSA clustering, we set the number of clusters used in the sampling for the predictions to vary between [16, 32, 64, 128, 256, 512, 1024, 5120] [5]. We take 13 samples per clustering threshold, resulting in 104 predictions per target in total (compared to the 50 used previously). In total, 145 and 154 out of 155 structures could be predicted using strategies 1 and 2, respectively. The number of resulting structures sampled is 14177 and 16007 for strategies 1 and 2, respectively. The failures are due to memory issues (NVIDIA A100 GPUs with 40 GB RAM were used).

We evaluate the predictions with the TM-score from TM-align, this time taking the maximum score to account for possible size differences between the native structures representing the different conformations (extracted matching regions) and the full genetic sequences used for prediction.

## Availability

Cfold is available for local installation here: https://github.com/patrickbryant1/Cfold/tree/master and as a Google Colab notebook here: https://colab.research.google.com/github/patrickbryant1/Cfold/blob/master/Cfold.ipynb

The data used to train the network and PDB files of the predictions used to generate the figures are available through: https://zenodo.org/record/8334555

## Acknowledgements

This study was supported by the European Commission (ERC CoG 772230 “ScaleCell”), MATH+ excellence cluster (AA1-6, AA1-10), Deutsche Forschungsgemeinschaft (SFB 1114/C03). Computational resources were obtained from ZIH (SCADS) at TU Dresden with project id p_scads_protein_na.

Blender 3D and Pymol were used to visualise the protein structures. We also thank DeepMind for making the AlphaFold2 code available and Atharva Kelkar for reading the manuscript.

